## Supplemental text and figures for "Effective mechanical potential of cell–cell interaction explains three-dimensional morphologies during early embryogenesis"

Koyama et al.

#### Contents

1. **Particle-based cell model**
2. **Acquisition of nuclear tracking data from mouse embryos**
3. **Handling of divided and dead/lost cells in nuclear tracking data**

   3-1. Handling of divided cells

   3-2. Handling of dead/lost cells
4. **Procedures for inferring effective forces of cell–cell interactions**

   4-1. Comparison of cell/particle positions *in vivo* vs. *in silico*

   4-2. Cost function of XYZ-coordinate errors

   4-3. Cost function of cell–cell interaction force

4-3-1. Definition of cut-off distance

4-3-2. Definition of cost function

4-3-3. Setting of coefficient of cost function

4-3-4. Setting of distance-dependent weight

4-3-5. Other possible cost functions

4-4. Minimization of cost function

4-4-1. Minimization under conditions with spatial constraints

**5. Data analysis of inferred effective forces of cell–cell interaction**

5-1. Generation of distance–force plot data

5-2. Estimation of distance–force curves

5-3. Calculation of distance–potential curves

5-4. Properties of distance–potential curves

**6. Simulation using distance–force curves**

6-1. Simulation based on distance–force curves derived from *in vivo* systems

6-2. Simulation data generation for systematic validation of inference method

6-2-1. Force fluctuation and its persistency

6-2-2. Simulations under various setting

**7. Experimental materials and methods**

7-1. Mouse embryos

7-2. Microscopic live imaging

**8. References**

**9. Supplementary figures**

**10. List of videos**

**11. List of source data**

#### 1. Particle-based cell model

In our particle-based cell model, particles interact with each other. Defining the range of particle–particle interaction distances is essential for performing simulations. In particle-based models such as those used for molecular dynamics simulations [1,2], when a distance of a pair of particles is large, the forces between the particles can become sufficiently small to be negligible. Thus, such interactions are non-effective on the behaviors of the systems. To exclude the non-effective interactions, a cut-off distance for particle–particle interactions is typically set: when a distance of each paired particles is larger than a threshold value, the force between the paired particles is set to be zero. In the case of cells, because cells can interact each other through various ways accompanying long protrusive structures, substrates, etc., it is technically difficult to identify all these interactions by microscopic imaging [3–7]. Therefore, in this study, we assumed that effective interaction distances were ~3-fold larger than the diameters of "cell bodies" which do not include process-portions of cells such as pseudopodia. Importantly, this cut-off distance did not substantially affect the results of inference of effective forces as shown later (Section 4-3-4). The diameters of cell bodies were estimated using the Voronoi tessellation.

The equation of particle motion is defined as described in the main text (Fig. 1B): $V_{C|p} = F_{C|p} / \gamma$. Due to the low Reynolds number in the case of cellular-level phenomena, the inertial force is negligible, in contrast to the simulations for molecular motions. Thermal fluctuations or noises of cellular activities are included as follows:

$$V_{C|p} = F_{C|p} / \gamma + \zeta, \qquad \text{(Eq. S1)},$$

where $\zeta$ is the fluctuation or noise. This is the equation of the motion of each particle. Because it is difficult to experimentally measure the value of $\gamma$ and its spatiotemporal distributions, we assumed the simplest situation, i.e., $\gamma$ is constant (=1.0). In addition, $\zeta$ was assumed to be negligible (=0), except for some cases as mentioned in Section 6. In the case of the relative velocity-dependent model shown in Fig. S8, the equation of particle motion was previously reported [8,9] as follows:

$$F_{C|p} = \sum_{m=1}^{N} [\{\omega_g(D_{pm})\gamma\}\{(V_{C|p} - V_{C|m}) \bullet e_{pm}\}e_{pm}], \qquad \text{(Eq. S1-2)},$$

where $m$ is the ID of particles interacting with the $p$th particle, $N$ is the total number of the interacting particles, $D_{pm}$ is the distance between the $p$th and $m$th particles, $\omega_g(D_{pm})$ is a distance-dependent weight for $\gamma$, and $e_{pm}$ is the unit vector from the XYZ-coordinates of $m$th to $p$th particles. $\omega_g(D_{pm})$ decays along $D_{pm}$ and becomes zero at the cut-off distance of cell–cell interactions. The decay function was described by a quadratic function [10].

#### 2. Acquisition of nuclear tracking data from mouse embryos

We utilized nuclear positions as the marker of cell positions, because the nuclear positions are widely available as mentioned in the main text. Although centroids of cells are candidate alternatives for cell positions, it is challenging to obtain centroid positions in three-dimensional tissues. Therefore, because our method uses nuclear positions, we expect that it could be used in a wide range of tissues and organisms. By utilizing confocal microscopy or other optical microscopy methods, nuclei labeled with fluorescence markers such as genetically encoded histone-GFP (green fluorescent protein) and chemical reagents such as Hoechst can be three-dimensionally imaged.

The confocal images obtained from the mouse embryos were subjected to the procedure for nuclear detection and tracking. Using the Imaris software (Oxford instruments/Bitplane, UK), the nuclei were automatically detected, followed by manual corrections. The nuclear tracking was also performed automatically followed by manual corrections. Through the manual corrections, the accuracies of the nuclear detection and tracking became essentially 100%, to the best of our judgment.

#### 3. Handling of divided and dead/lost cells in nuclear tracking data

To make the nuclear tracking data applicable to the procedures for inference of cell–cell interaction forces, we modified the tracking data of the *C. elegans* and mouse embryos. As described later in Section 4, force inference was performed for each time frame. For instance, when the effective mean forces were inferred at time frame = $t$ in Figure S1B, the nuclear movement data at $t$ and $t + \Delta t$ were used. In other words, a pair of time frames was considered at each time frame: in Figure S1B, there are three pairs, [$t$-2$\Delta t$, $t$-$\Delta t$], [$t$-$\Delta t$, $t$], and [$t$, $t$+$\Delta t$]. In cases that do not contain dividing and dead/lost cells, both cell numbers and cell IDs are identical between the paired time frames (Fig. S1B; [$t$-2$\Delta t$, $t$-$\Delta t$] and [$t$, $t$+$\Delta t$]), and this data format is applicable to our force inference method. By contrast, in cases containing dividing and dead/lost cells, neither the cell numbers nor the cell IDs are the same (Fig. S1B; [$t$-$\Delta t$, $t$]), and this data format is not applicable to our force inference method. To make the latter data format applicable to our force inference method, we modified the data to have the same cell numbers and cell IDs between the paired time frames as follows.

#### 3-1. Handling of divided cells

A mother cell and its daughter cells have different IDs (Fig. S1B; ID = 1 for a mother cell, and ID = 2 and 3 for two daughter cells). At time frame nearest to cell division, a mother cell was virtually assumed to be positioned at the centroid of its two daughter cells (Fig. S1B; *t*, ID = 1), and the daughter cells are excluded. Consequently, both the cell numbers and the cell IDs became the same in the paired time frames (Fig. S1B; [$t$-$\Delta t$, $t$]). On the other hand, at the next paired time frames, the virtually assumed-mother cell was excluded, and the daughter cells are included (Fig. S1B; [$t$, $t$+$\Delta t$]), resulting in the same cell numbers and cell IDs in the paired time frames. Divided cells appeared in the *C. elegans* and mouse embryos.

#### 3-2. Handling of dead/lost cells

Similar to the handling of divided cells above, dead and lost cells lead to different cell numbers and cell IDs in the paired time frames. To eliminate this inconsistency, these cells were excluded from the paired time frames. Dead and lost cells appeared in the *C. elegans* embryo.

### 4. Procedures for inferring effective forces of cell–cell interactions

To infer mean effective cell–cell interaction forces, the particle-based cell simulation model was fitted to the nuclear tracking data obtained from *in vivo* observations in the spirit of data assimilation. Data assimilation is a technique to solve an inverse problem for a simulation-based model but not a function-based model such as a linear regression [11,12]. In our method, the effective force is inferred for each cell–cell interaction for each time frame. Thus, a network of the interaction forces for each time frame is obtained (Fig. 1B-iii, 4C, and S2B), and simultaneously, the temporal evolution of forces of each cell–cell interaction is also obtained.

In this section, we explain the principles and procedures of the fitting method. Briefly, we systematically searched for effective mean forces of cell–cell interactions that could reproduce the *in* *vivo* nuclear mean velocities/movements (Fig. 1B). This search was performed for each pair of time frames (Section 4-1). The search process corresponds to the minimization of a cost function, which is defined as the differences in nuclear movements between *in vivo* and *in silico* (Section 4-2 and 4-4). A cut-off distance of cell–cell interactions was considered (Section 4-3-1), and this constraint was implemented in the cost function to be minimized (Sections 4-3-2, -3, and -4). A unique solution of a cell–cell interaction force was obtained through minimization of the cost function (Section 4-4).

#### 4-1. Comparison of cell/particle positions *in vivo* vs. *in silico*

From nuclear tracking data, as mentioned in Section 3, we generated sequential pairs of time frames that contain the same cell numbers and cell IDs and inferred the mean forces for each pair (Fig, S1B). In a pair of time frames (e.g. [*t, t+Δt*] in Fig. S1B), the mean velocities/movements of the nuclei were obtained from the nuclear tracking data. We defined that the nuclear positions in the first time frame (*t*) in the pair [*t, t+Δt*] as the initial particle positions in the simulations described below. In the particle-based cell model, mean cell–cell interaction forces were given, and a simulation was performed from *t* using the Euler method, where the time step was set to be the same as the time interval of the nuclear tracking data *Δt*. Note that the time interval of the nuclear tracking data was set to be small so that nuclear movements were almost smooth, which may be essential to perform the fitting. The resultant particle positions at time frame = *t+Δt* were compared with those of the nuclear tracking data, and the positional difference was evaluated. To minimize the difference, the mean cell–cell interaction forces were optimized. During this inference, we assumed the simplest situation, where $\gamma$ is constant (=1.0) and $\zeta$ is negligible (=0).

#### 4-2. Cost function of XYZ-coordinate errors

The positional difference between the nuclei (*in vivo*) and the particles (*in silico*) mentioned above is defined as follows. Because the values of the time interval *Δt* can differ among nuclear tracking data or even among different time frames of the same tracking data, we considered that the cost function to be minimized should not depend on the differences in the value of the time interval.

Here we consider a pair of time frames [*t, t+Δt*] in Figure S1B. The XYZ coordinate of the *p*th nuclei at time frame *t* and *t+Δt* are defined as $r_{t,p}{}^{\mathrm{ref}}(0) = (x_{t,p}{}^{\mathrm{ref}}(0), y_{t,p}{}^{\mathrm{ref}}(0), z_{t,p}{}^{\mathrm{ref}}(0))$ and

$r_{t,p}^{\text{ref}}(\Delta t) = (x_{t,p}^{\text{ref}}(\Delta t), y_{t,p}^{\text{ref}}(\Delta t), z_{t,p}^{\text{ref}}(\Delta t))$, respectively, where ref means reference (Fig. S1B). The coordinates of corresponding particles in simulations are defined as $r_{t,p}(0) = (x_{t,p}(0), y_{t,p}(0), z_{t,p}(0))$ and $r_{t,p}(\Delta t) = (x_{t,p}(\Delta t), y_{t,p}(\Delta t), z_{t,p}(\Delta t))$, respectively. The distances between a nucleus and a particle at $t$ and $t+\Delta t$ are defined $s_{t,p}(0)$ and $s_{t,p}(\Delta t)$ where $s_{t,p}(0) = |r_{t,p}(0) - r_{t,p}^{\text{ref}}(0)|$ and $s_{t,p}(\Delta t) = |r_{t,p}(\Delta t) - r_{t,p}^{\text{ref}}(\Delta t)|$, respectively. Note that $s_{t,p}(0) = 0$, because the initial particle positions of simulations are equivalent to those from nuclear tracking data, as explained in Section 4-1.

During a sufficiently short time period $\tau$, the movement trajectories of both the nucleus and the particle can be approximated as linear. Thus, if the values of $\Delta t$ are small enough ($\Delta t \le \tau$), the distance between the nucleus and the particle would increase linearly during the time period $\Delta t$. Because $s_{t,p}$ would increase linearly $(s_{t,p}(\Delta t) \propto \Delta t)$, $s_{t,p}(\Delta t) / \Delta t$ is constant and does not depend on $\Delta t$ during time period $\tau$. Note that $s_{t,p}(\Delta t) / \Delta t$ corresponds to the increasing rate of $s_{t,p}$. For convenience, we used $\{s_{t,p}(\Delta t) / \Delta t\}^2$, which is also independent to $\Delta t$. Then, we defined the positional differences produced during $\Delta t$ as $\{s_{t,p}(\Delta t) / \Delta t\}^2 \Delta t$. Under this definition, the values of the positional differences produced during the same time period do not depend on the time interval of the nuclear tracking data.

The summation of the positional differences of all particles through all time frames is written as follows:

$$\begin{aligned} G_{xyz} &= \sum_{t=1}^{T} \sum_{p=1}^{P(t)} \left\{ \frac{s_{t,p}(\Delta t)}{\Delta t} \right\}^2 \Delta t \\ &= \sum_{t=1}^{T} \sum_{p=1}^{P(t)} \left\{ \frac{\left| r_{t,p}(\Delta t) - r_{t,p}^{\text{ref}}(\Delta t) \right|}{\Delta t} \right\}^2 \Delta t = \sum_{t=1}^{T} \sum_{p=1}^{P(t)} \frac{\left\{ r_{t,p}(\Delta t) - r_{t,p}^{\text{ref}}(\Delta t) \right\}^2}{\Delta t} \end{aligned} \quad \text{Eq. S2,}$$

where $p$ is the particle ID, and $P(t)$ is the total number of particles at $t$, which varies by $t$ due to cell division etc. Here we defined $r_p^{\mathrm{ref}}(t) = (x_p^{\mathrm{ref}}(t), y_p^{\mathrm{ref}}(t), z_p^{\mathrm{ref}}(t)) = (x_{t,p}^{\mathrm{ref}}(\Delta t), y_{t,p}^{\mathrm{ref}}(\Delta t), z_{t,p}^{\mathrm{ref}}(\Delta t))$ and $r_p(t) = (x_p(t), y_p(t), z_p(t)) = (x_{t,p}(\Delta t), y_{t,p}(\Delta t), z_{t,p}(\Delta t))$ for convenience, and obtained

$$G_{xyz} = \sum_{t=1}^{T}\sum_{p=1}^{P(t)} \frac{\left\{r_p(t) - r_p^{\mathrm{ref}}(t)\right\}^2}{\Delta t}$$
$$= \sum_{t=1}^{T}\sum_{p=1}^{P(t)} \frac{\left\{x_p(t) - x_p^{\mathrm{ref}}(t)\right\}^2 + \left\{y_p(t) - y_p^{\mathrm{ref}}(t)\right\}^2 + \left\{z_p(t) - z_p^{\mathrm{ref}}(t)\right\}^2}{\Delta t} \qquad \text{Eq. S3}$$

, which corresponds to Equation 2 in the main text. This cost function was minimized. In addition, by defining $s_p(t) = s_{t,p}(\Delta t)$, this equation can be interpreted as $G_{xyz} \simeq \sum_{p=1}^{\bar{P}} \int_1^T \left\{ \frac{ds_p(t)}{dt} \right\}^2 dt$, where the total number of particles is $\bar{P}$.

#### 4-3. Cost function of cell–cell interaction force

In the particle-based cell model, we defined a cut-off distance for cell–cell interactions (Section 1). Consistency between inferred effective forces and the cut-off distance should be maintained. Considering a cut-off distance is equivalent to making an assumption that the effective forces are zero at longer distances than the cut-off distance (Fig. S1A-ii). When we infer the forces, cell–cell interactions whose distances are longer than the cut-off distance are neglected; hence, the forces of the interactions automatically become zero (Fig. S1A-ii, right side of the red broken line). On the other hand, in the case that the distances of cell–cell interactions are slightly shorter than the cut-off distance, the forces should be nearly zero and not have large absolute values (Fig. S1A-ii; red crosses that have large absolute values should not be permitted). Otherwise, a discontinuity in the relationship between the forces and the particle–particle distances would be produced at the cut-off distance, which is not physically reasonable (Fig. S1A-ii), because the values of the effective forces would change abruptly at the cut-off distance. To achieve the correct profile of the effective forces mentioned above, we assumed that a constraint for force values was incorporated into the function to be minimized during the inference. In the following sections, we describe the definition of the cut-off distance, as well as the definition and actual setting of the function to be minimized.

#### 4-3-1. Definition of cut-off distance

We initially assumed that effective cell–cell interaction distances were ~ 3-fold greater than the diameters of the cell bodies, i.e., the non-process portion of cells. However, this cut-off distance did not affect the values of inferred forces as shown later (Section 4-3-4). To estimate the diameters from the nuclear images, we utilized the Voronoi tessellation (Fig. S1A-i). In three-dimensional systems, including the *C. elegans* and mouse embryos, three-dimensional tessellations were used. Neighboring cells were defined using the Voronoi tessellation, and the distances between pairs of neighboring cells were calculated. In the mouse embryos, we calculated an averaged value of the distances of all cell–cell interactions across all time frames for each embryo, and defined the value as the characteristic diameter of the cells for each embryo. In the *C. elegans* embryo, the characteristic diameter was defined for each time frame, because the embryo experiences multiple cell divisions that cause substantial changes in the diameters.

#### 4-3-2. Definition of cost function

We defined a cost function for cell–cell interaction forces in order to diminish the discontinuity in the relationship between the forces and the distances at the cut-off distance. We assumed that the effective forces ($F_i$) of cell–cell interactions should approach zero when the cell–cell distances ($D$) approach the cut-off distance. Thus, we considered $|F_i|$ or $F_i^2$ as the cost around the cut-off distance. Furthermore, because discontinuities of the relationship should not appear at any distance, we set the cost to be distance-dependent and to gradually increase around the cut-off distance. To satisfy this, we set the cost as $\Psi(D)\ F_i^2$, where $\Psi(D)$ is a distance-dependent weight that increases gradually toward the cut-off distance and eventually becomes extremely large, as shown in Figure S1A-iii. We defined that

$$\psi(D) = \alpha^{(D-D_c)/D_c} \qquad \text{Eq. S4,}$$

where $\alpha \geq 1$ and $D_c$ is the diameter of the cell bodies estimated by the Voronoi tessellation. If $\Psi(D)$ was set to be constant, inferred effective forces became far from zero even at long distant regions as shown later. The setting of the value of $\alpha$ will be discussed later.

We also considered that the cost function should not depend on the differences in the value of the time interval $\Delta t$ for the same reason mentioned in Section 4-2. Hence, we defined the cost produced during $\Delta t$ is $\Psi(D)\ F_i^2 \Delta t$. Under this definition, the values of the costs produced during the same time period $\tau$ do not depend on the time interval $\Delta t$ of the nuclear tracking data, according to the same logic explained in Section 4-2. Then, we obtained the total cost as follows

$$g_{F_0} = \sum_{t=1}^{T} \sum_{i=1}^{I(t)} \psi(D_i) F_i^2 \Delta t \qquad \text{Eq. S5,}$$

where time frame ($t$) is from 1 to $T$, and the cell–cell interaction ID ($i$) defined in each time frame is from 1 to $I(t)$. This equation can be interpreted as $g_{F_0} \simeq \sum_{i=1}^{\bar{I}} \int_1^T \psi(D_i(t)) F_i^2 dt$, where the cell–cell interaction ID ($i$) is defined for each pair of particles and is conserved across all time frames. The total number of pairs is $\bar{I}$. In addition, this term can be considered as the prior in the Bayesian inference. Therefore, our minimization problem is a fitting of a simulation-based model (i.e. data assimilation) combined with the Bayesian approach (i.e. with a prior). Distance-dependent constraints other than Equation S5 were assessed, but they did not improve the inference results as shown later.

##### 4-3-3. Setting of coefficient of cost function

We defined the summation of the Equations S3 and S5 as the cost function to be minimized, resulting in the Equation 3 in the main text. When we added the cost of the effective forces of cell–cell interactions $g_{F_0}$ (Section 4-3-2) to the cost of the XYZ coordinate $G_{xyz}$ (Section 4-2), we introduced a coefficient $\omega_{F_0}$ related to $g_{F_0}$ as follows,

$$G = G_{xyz} + G_{F_0} \quad (G_{F_0} = \omega_{F_0} g_{F_0}) \qquad \text{Eq. S6.}$$

$\omega_{F_0}$ determines the relative contribution of $g_{F_0}$ to $G$. Importantly, in the absence of the second term, this minimization problem exhibited indefiniteness (i.e. a unique solution could not be determined) and over-fitting also occurred. The second term is considered as the prior in the Bayesian inference, and, in general, a unique solution could be ensured by introducing such a prior [11]. $\omega_{F_0}$ was set to be small to ensure that the value of $G_{xyz}$ does not show large values after minimization of $G$. Otherwise, it is expected that inferred forces cannot reproduce the *in vivo* nuclear positions with high accuracy. In addition, because the dimension of $g_{F_0}$ is different from that of $G_{xyz}$, $\omega_{F_0}$ is defined in such a manner that it causes $G_{xyz}$ and $G_{F_0}$ to have the same dimension. We determined the value of $\omega_{F_0}$ as follows.

###### 4-3-3-1. Definition of accuracy

The value of the coefficient ( $\omega_{F_0}$ ) was set so that the particle positions in the simulations exhibited ~98% consistency (i.e. accuracy) with the *in vivo* nuclear positions. Here, we first showed the definition of the accuracy of the reproduction of particle positions calculated by using $G_{xyz}$ , and then, showed how the value of $\omega_{F_0}$ was set.

If particle positions (*in silico*) obtained from inferred forces of cell–cell interactions are absolutely consistent with nuclear positions (*in vivo*), $G_{xyz}$ becomes 0, and the accuracy was defined as 100 %. This situation coincides with $x_p(t+\Delta t) = x_p^{\mathrm{ref}}(t+\Delta t)$ , $y_p(t+\Delta t) = y_p^{\mathrm{ref}}(t+\Delta t)$ , and $z_p(t+\Delta t) = z_p^{\mathrm{ref}}(t+\Delta t)$ in Equations S3. When particle positions show no movements from *t* to (*t* + *Δt*), which can be yielded under all cell–cell interaction forces = 0, the cost function $G_{xyz}$ will have a large value

$$G_{xyz}{}^{0} = \sum_{t=1}^{T}\sum_{p=1}^{P(t)} \frac{(x_p^{\mathrm{ref}}(t) - x_p^{\mathrm{ref}}(t+\Delta t))^2 + (y_p^{\mathrm{ref}}(t) - y_p^{\mathrm{ref}}(t+\Delta t))^2 + (z_p^{\mathrm{ref}}(t) - z_p^{\mathrm{ref}}(t+\Delta t))^2}{\Delta t}$$

$= \sum_{t=1}^{T}\sum_{p=1}^{P(t)} \frac{(s_p{}^{0})^2}{\Delta t}$ . When $G_{xyz}$ is equal to $G_{xyz}{}^{0}$ , we defined that the accuracy was 0 %; the superscripts of $G_{xyz}{}^{0}$ and $s_p{}^{0}$ means 0 %. Here, we defined $G_{xyz}{}^{Q}$ as follows;

$$G_{xyz}{}^{Q} = G_{xyz}{}^{0}\left(\frac{100-Q}{100}\right)^2 \qquad \text{Eq. S7}$$

When $G_{xyz}$ is equal to $G_{xyz}{}^{Q}$ , the accuracy was defined as $Q$ %.

Usually, 100% accuracy was not achieved even under any values of cell–cell interaction forces. This is because nuclear tracking data contain various kinds of noise or observation errors. For instance, when the centroid of tissues can be moved due to a drift, such translational motion cannot be reproduced by the particle-based cell model where $\gamma$ (the coefficient of the viscous drag force) is constant. Rotational motions of tissues were also detected in floating tissues. As far as our examinations showed, simulations of the particle-based cell model cannot cause translational or rotation motions of a particle cluster if $\gamma$ is constant, probably because external forces are required to generate these motions. Actual nuclear tracking data contained these motions because the embryos not adhered to substrates were easily moved and rotated by slight flows of the surrounding medium. Many other factors can also cause these motions, although we cannot enumerate all of them. Microscopic systems are affected by slight changes in room temperature, leading to movements of the sample stages of the systems. Nuclear detection errors also yield these motions.

##### 4-3-3-2. Canceling of translational and rotational motions of nuclei

Because the translation and rotational motions varied significantly among time frames in each set of tracking data and also among tracking data, it was necessary to cancel these motions. Otherwise, the accuracy mentioned in Section 4-3-3-1 could not be fairly evaluated. Therefore, we modified the nuclear positions in each pair of time frames. Note that these modifications did not eventually affect the inference results: under the same value of $\omega_{F_0}$, the values of the inferred forces became the same between the cases with and without modifications (data not shown). In other words, this canceling process is solely used for fairly calculating the accuracy.

The translational motions were easily canceled: we translationally moved the nuclei in one of the two time frames ($t$ and $t + \Delta t$) of each pair so as to give the same position of the centroid as that in another time frame.

After the correction of the translation motions, the rotational motions between the two time frames ($t$ and $t + \Delta t$) of each pair were eliminated by three-dimensionally rotating the positions of the nuclei at $t + \Delta t$ as follows. We defined that the centroid of the nuclei as the origin of the XYZ coordinates, allowing us to write the position vectors of the nuclei for the two time frames as $\vec{r}_i(t) = (x_i(t), y_i(t), z_i(t))$, where $i$ is the $i$th nucleus. The rotational motion of each nucleus between the two time frames may be evaluated by the cross product $\vec{L}_i = \vec{r}_i(t) \times \left\{ \vec{r}_i(t+\Delta t) - \vec{r}_i(t) \right\}$, which is analogous to angular momentum. Thus, the total rotational motions of all nuclei may be evaluated by $\sum_{i=1}^{I(t)} |\vec{L}_i|^2$, and we three-dimensionally rotated the positions of the nuclei so that this cost function was minimized. A three-dimensional rotation is generally achieved by the following calculation;

$$\begin{pmatrix} x_i' \\ y_i' \\ z_i' \\ 1 \end{pmatrix} = \begin{pmatrix} \cos\phi\cos\theta & \cos\phi\sin\theta\sin\psi - \sin\phi\cos\psi & \cos\phi\sin\theta\cos\psi + \sin\phi\sin\psi & 0 \\ \sin\phi\cos\theta & \sin\phi\sin\theta\sin\psi + \cos\phi\cos\psi & \sin\phi\sin\theta\cos\psi - \cos\phi\sin\psi & 0 \\ -\sin\theta & \cos\theta\sin\psi & \cos\theta\cos\psi & 0 \\ 0 & 0 & 0 & 1 \end{pmatrix} \begin{pmatrix} x_i \\ y_i \\ z_i \\ 1 \end{pmatrix}$$

, where $\phi$, $\theta$, and $\psi$ are the angles expressing the three-dimensional rotations, and $x_i'$, $y_i'$ and $z_i'$ are the resultant coordinates. Thus, to minimize the cost function, we searched for the values of these angles at $t + \Delta t$. The minimization was achieved using a Monte Carlo algorithm.

After modification of these motions, we were able to find cell–cell interaction forces that achieved almost 100% accuracy by minimizing $G_{xyz}$ in all cases of the present study, indicating that the definition of the cost function and the subsequent Monte Carlo algorithm worked sufficiently well to cancel the rotational motions. Note that minimization of $G_{xyz}$ would cause over-fitting and did not yield a unique solution for the cell–cell interaction forces. By introducing $G_{F_0}$, a unique solution would be obtained, as described below. In addition, to achieve 100% accuracy, a longer cut-off distance may be required. If the cut-off distance was 3 times the cell diameter, 100% accuracy was always achieved. However, when effective cell–cell interactions were only assumed between neighboring cells determined by the Voronoi tessellation, 100% accuracy was usually not achieved, suggesting that cut-off distance comparable to the cell diameter may not be sufficient.

###### 4-3-3-3. Determination of the value of $\omega_{F_0}$

After correcting for translational and rotational motions, we performed minimization of $G = G_{xyz} + G_{F_0}$ ($G_{F_0} = \omega_{F_0} g_{F_0}$) under various values of $\omega_{F_0}$ with $\Psi(D)$ = 1 (const.), and searched for the values satisfying that the accuracy was 98 ± 0.2 %. The value of $\omega_{F_0}$ was determined for each set of nuclear tracking data, and the same value was applied through all time frames for each dataset. Importantly, once the value of $\omega_{F_0}$ was determined, the values of inferred forces were not affected by modification of translational and rotational motions (data not shown). Thus, the modification of these motions is only meaningful for the determination of $\omega_{F_0}$ but not for the inference results. Additionally, when we set the accuracy to be achieved to 99.5 % instead of 98 %, the inference results were not significantly affected, and essentially similar distance–force curves were obtained (data not shown). Furthermore, for these accuracies, a unique solution could be obtained as shown later.

##### 4-3-4. Setting of distance-dependent weight

The value of the distance-dependent weight $\alpha$ in $\Psi(D)$ ($= \alpha^{(D-D_c)/D_c}$) was determined by using simulation data of particle tracking under pre-given potentials (Fig. S2A, Lenard–Jones and Freehand potentials), and then, the value was commonly applied to all nuclear tracking data shown in this study. In other words, we searched for a good value of $\alpha$ that yielded cell–cell interaction force values consistent with the various pre-given potentials.

Under this value of *α,* the values of inferred effective forces were essentially the same under different cut-off distances (Fig. S6). In the cases where the cut-off distances were greater than the 3-fold diameter of cell bodies (Fig. S6, 3.2 and 4.0-fold), the values of the inferred effective forces were almost identical among the different cut-off distances in the *C. elegans* embryos. Shorter cut-off distances caused increased differences in the values of the inferred effective forces (Fig. S6, 2.0 and 2.5-fold). Therefore, our inference results do not depend on the cut-off distances if the cut-off distances were set to be greater than 3-fold.

##### 4-3-4-1. Simulation data of particle tracking under various potentials

We generated particle tracking data by performing simulations under various potentials. We first utilized the Lenard–Jones (LJ) potential with various force scales. The LJ potential is written as follows; $W(D)=\kappa\left\{\left(\frac{\sigma}{D}\right)^{12}-\left(\frac{\sigma}{D}\right)^{6}\right\}$, where the potential *W* is described as a function of the particle–particle distance *D*. The first and second terms correspond to the repulsive and attractive forces, respectively. $\kappa$ determines the potential and force scales. $\sigma$ almost corresponds to the diameter of the particle. Then, we can express this distance–potential curve as a distance–force curve as follows; $F(D)=\frac{\kappa}{\sigma}\left\{12\left(\frac{\sigma}{D}\right)^{13}-6\left(\frac{\sigma}{D}\right)^{7}\right\}$. Basically, the profiles of the attractive and repulsive terms are similar to those in Fig. 1A-iii. We generated the LJ potential-based distance–force curves with various $\kappa$ values, and then performed simulations, and finally obtained particle tracking data (Fig. S2A and E). In the simulations, we set $\zeta = 0$ and $\gamma = 1$. We set various initial configurations of the particles and various values of the time step of the iteration of the simulations and of the sampling time interval. We also defined a freehand distance–force curve called FH potential (Fig. S2A). Similar to the case of the LJ potential, we generated distance–force curves based on the FH potentials with different force scales followed by simulations, and then obtained particle tracking data (Fig. S2A and E). We also defined distance–force curves by translating the LJ and FH potentials along the direction of the distance (data not shown). As a negative control, particle tracking data of random walks were also generated (Fig. S2A). We provide these distance–force curves as text files (Source Data). Simulation data related to Figure 2, 3, and S4-5 were generated as described in Section 6-2.

##### 4-3-4-2. Determination of $\alpha$

For each particle tracking data described in Section 4-3-4-1, $\omega_{F_0}$ was determined by the procedures shown in Section 4-3-3-3. Then, we searched for the values of $\alpha$ that can yield inference results which are well consistent with the original distance–force curves based on the LJ and FH potentials.

Figure S2C exemplifies a few inference results obtained under good values of $\alpha$. The values of distance and inferred force for each cell–cell interaction for each time frame were plotted, and heat maps were generated from the frequencies of the data points (left panels). For instance, many data points were plotted around the red regions, whereas few data points were plotted around the black regions. In the white regions, no data points were plotted. In the right panels of Figure S2C, the original distance–force curves (orange) and the inferred distance–forces curves (yellow) estimated from the plot data are overlaid on the left panels. The plots were almost distributed around the original distance–force curves. These inferred curves were nearly consistent with the original curves in the cases of the LJ and FH potentials. By contrast, in the case of the random walk, the plots were widely distributed, and no clear distance–force curves were obtained (yellow dots).

Some examples of the effect of $\alpha$ values on inference results are shown in Figure S2D. When $\alpha$ was 1.0, the distance–force curves were flatter than the original curves of the LJ and FH potentials. When the value of $\alpha$ was increased (1, 5, 30, 100, 300), the inferred distance–force curves showed clear peaks at similar distances of the original curves, and around $\alpha$ = 100, the inferred curves approached the original curves quite closely. When $\alpha$ =100, the scales of forces in the original curves were almost identical to the inferred curves, as shown in Figure S2E: the values of the forces in the original curves at the peaks were 0.03 (red), 0.01 (green), 0.003 (light blue), and 0.001 (purple), and the values in the inferred curves at the peaks were consistent with or slightly smaller than those values. We also applied this value of $\alpha$ to other distance–force curves that were obtained by translating the LJ and FH potentials along the direction of the distance (Section 4-3-4-1), and found that inferred curves closely reflected the translations (data not shown). In addition, under this value of $\alpha$, no clear distance–force curves were obtained from the random walk data. Thus, we can expect that if a distance–force curve exists in a system of interest, the curve will be almost correctly inferred, whereas if no distance–force curve exists, no clear curve will be obtained. Finally, this value of $\alpha$ was applied to nuclear tracking data obtained *in vivo*.

##### 4-3-5. Other possible cost functions

We also tested other cost functions in order to further improve inference results. Instead of the $F_i^2$ of the cost function $g_{F_0} = \sum_{t=1}^{T}\sum_{i=1}^{I(t)} \psi(D_i) F_i^2 \Delta t$, we tested $|F_i|$ as previously described in Section 4-3-2. Moreover, we tried to implement a cost for the temporal smoothness of forces for each cell–cell interaction by $g_{F_1} \simeq \sum_{i=1}^{\bar{I}} \int_1^T \psi_{F_1}(D_i(t)) \left( \frac{dF_i}{dt} \right)^2 dt$ or by $g_{F_2} \simeq \sum_{i=1}^{\bar{I}} \int_1^T \psi_{F_2}(D_i(t)) \left( \frac{d^2 F_i}{dt^2} \right)^2 dt$. $g_{F_0}$, $g_{F_1}$, and $g_{F_2}$ contain costs derived from the zero-, first-, and second-order differentials of the forces, respectively. We tested various combinations of these costs, and found that some combinations improved the inference results for very specific particle tracking data whereas they made the results worse for other tracking data (data not shown). Through these evaluations, we concluded that the cost $g_{F_0}$ was most widely applicable to various distance–force curves, though we do not rule out the possibility that better cost functions could be found in the future.

#### 4-4. Minimization of cost function

To minimize the cost function $G = G_{xyz} + G_{F_0}$, we numerically solved by using the conjugate gradient method whose algorithm is described in *Numerical Recipes in C* (Cambridge University Press). Basically, values of particle–particle interaction forces are repeatedly improved in order to minimize the cost function. Initial values of particle–particle interaction forces are set to 0, but we confirmed that essentially identical values of the forces were obtained even if the minimization was started from various non-extreme initial values (Fig. 4D and S6), suggesting that a unique solution was obtained. On the other hand, in the absence of $G_{F_0}$, no unique solution was obtained, leading to overfitting. In Fig. 6B-C, the initial values of the forces were differently set among the data points in the right panels: different uniform random numbers were provided ([from -0.02 to 0.02] in B and [from 0.003 to 0.003] in C).

During force inference, we assumed that $\gamma$ is constant (=1.0) and $\zeta$ is negligible (=0), as mentioned in Section 4-1. The dimension of particle velocity $V$ was set to be μm/min for convenience, which is typical dimension in analyses of living cells. Because we did not measure the value of $\gamma$ in living tissues, we cannot determine the value of the effective force $F_{C|p}$ ($=\gamma V_{C|p}$) with a specific dimension. However, under the above assumption, 1 arbitrary unit (A.U.) of $F$ coincides to a force which can move a particle with a velocity of 1 μm/min in tissues. Therefore, we can evaluate the effect of forces on the movements of cells in tissues. We can also say that the effects of forces on cellular movements can be compared among different tissues, even if those tissues have different values of $\gamma$. In other words, we can compare the values of $E_{C|p} = F_{C|p} / \gamma$ among different tissues, where $E_{C|p}$ is an index of the effect of forces on cellular movements.

In the case of the relative velocity-dependent model (Fig. S8 and Eq. S2-1), we used $V_{C|m}{}^{\mathrm{ref}}$ instead of $V_{C|m}$ in Eq. S2-1 as follows; $F_{C|p} = \sum_{m=1}^{N} [\{\omega_{\mathrm{g}}(D_{pm})\gamma\}\{(V_{C|p} - V_{C|m}^{\mathrm{ref}}) \bullet e_{pm}\} e_{pm}]$, where $V_{C|m}{}^{\mathrm{ref}}$ is the velocity of $m$th particle obtained from the cell tracking data. By introducing this approximation, Eq. S2-1 becomes a ternary system of linear equations for each particle; the three variables in this system are the XYZ components of $V_{C|p}$. Consequently, we can calculate $V_{C|p}$ by providing $F_{C|p}$, and thus, we can apply the cost function ($G$) for this model.

##### 4-4-1. Minimization under conditions with spatial constraints

In Figure 3, and S4-5, spatial constraints were included in the simulations. Similarly, the *C. elegans* and mouse embryos possess spatial constraints such as eggshells and zona pellucida. During the minimization of the cost function, these constraints were not explicitly considered.

#### 5. Data analysis of inferred effective forces of cell–cell interaction

Effective forces for each cell–cell interaction were inferred for each time frame by the procedures described in Section 4. Here, we present subsequent procedures for data analyses of the forces, which include plotting of the forces in distance–force space, estimation of distance–force curves, and calculation of distance–potential curves.

#### 5-1. Generation of distance-force plot data

Inferred effective forces were plotted in distance–force space. Because there were a lot of data points, we utilized a heat map to visualize where the data points were densely distributed in space. To make a heat map, we first prepared an image with 128×128 pixels, whose x- and y-axes corresponded to distance and force, respectively. Then, we set the minimum and maximum values of the distance on the image. In the case of Figure 4E, the values were -5 and 30 μm, respectively. The minimum and maximum values of the force were set to be -0.1 and 0.1 A.U., respectively. Then, the space was divided into the 64×64 square regions composed of 2×2 pixels, where the x- and y-widths of each region were {30-(-5)} / 64 = 0.55 μm and {0.1-(-0.1)} / 64 = 0.0031A.U., respectively. The outer distance–force space of the image was also divided into regions with these x- and y- widths. Then, the data points were allocated to one of the 64×64 square regions, or the regions in the outer space. Then, the counts for each square region were computed. We considered 64 columnar regions along the distance, and the mean values of the counts of the square regions were calculated for each columnar region. Note that the regions with counts = 0 were excluded from the calculation. The mean count for each 64-column region was defined as the frequency index (FI) = 1. Finally, the pixels were colored according to FI. The square regions were set to be 64×64 (2×2 pixels) for Figure 4E; and 128×128 (1×1 pixels) for Figure 2A and S2C. The minimum and maximum values of the distances and forces are shown in the Figures, and the x- and y- widths of each region differed according to the minimum and maximum values. These procedures were implemented in C using OpenCV.

#### 5-2. Estimation of distance–force curves

To estimate distance–force curves, we focused on the 64 columns along the distance in the case of Figure 4E, which corresponded to the x-width of the square regions shown in Section 5-1. We calculated the average values of the forces of data points plotted in each width of the 64 columns. The values of the distances of the data points were also averaged. The resultant values of the forces and distances were plotted as yellow boxes on the distance–force image defined in Section 5-1 (Fig. 4E etc.). The resultant values were then smoothed along the distance by averaging the values of the three serial data, and distance–force curves were obtained (Fig. 4F etc.). This smoothing was not applied to simulation data (Fig. 2A etc.).

To estimate distance–force curves of interactions of the outer–outer cells, the inner–inner cells, and the outer–inner cells in the compaction stage, in Figure S10, each cell type was manually identified in microscopic images, and distance–force plot data were generated for each interaction. The discrimination between outer and inner cells in the compaction stage embryos was occasionally difficult. However, such difficulty did not frequently occur, thus we believe that overall trends of the distance–force curves were not significantly affected by these errors.

In Fig. S10, the curves of the inner–inner and the outer–outer cells show different profiles, which may be consistent with previous works suggesting mechanical differences between the outer and inner cells [13]. The potential minimum of the inner–inner cells is lowest, which is consistent with the differential adhesive hypothesis where cells with strongest cell–cell adhesion should assemble [14]. The distance of the potential minimum of the outer–outer cells is longest, indicating longest-range of interactions among the three interactions. This may reflect the sealing of the embryonic surface by the outer cells as previously described [15]. The potential minimum of the inner–outer cells is lower than that of the outer–outer cells may reflect the reduced surface tension of the inner–outer cell boundaries as shown previously [13], which also lead to the sealing of the embryonic surface by the outer cells.

#### 5-3. Calculation of distance–potential curves

To calculate distance–potential curves, we mathematically integrated the distance–force curves obtained in Section 5-2 from the cut-off distance to distance = 0 as follows. Because the intervals between the data points in the distance–force curves were too wide to get smooth distance–potential curves, we linearly interpolated between the intervals, and then added new data points on the interpolated lines within a narrow interval. For the newly added data points, as well as the original ones, we performed integration from the cut-off distance, resulting in generation of distance–potential curves (Fig. 4F etc.).

#### 5-4. Properties of distance–potential curves

We analyzed the properties of the inferred distance–potential curves as follows.

##### 5-4-1. Slope of distance–potential curves

In Fig. 7C, we defined an index for evaluating long-range interactions. First, we searched for the distances at potential minima in the DP curves. Then, distances providing a given % of the potential minima were measured (Fig. 7C, right panel, the given % = 10%; Fig. S14A; Fig. S14B, the given % = 1). We used the distances relative to the distances at the potential minima as the index. If this relative distance is large, long-range attractive forces are exerted. In Figure 7C, 8B, and 8C, Mann–Whitney–Wilcoxon tests were performed implemented in R software as the "wilcox.exact" function.

#### 6. Simulation using distance–force curves

We performed simulations using distance–force curves. The simulations were performed by the Euler method written in C. The noise or fluctuation $\zeta$ in the Equation S1 was essentially set to 0. The fluctuations in simulation data for validating the inference method in Figure 2 and 3 are mentioned later.

##### 6-1. Simulation based on distance–force curves derived from *in vivo* systems

As in systems composed of larger colloidal particles which hardly reach a thermodynamically equilibrium state [16], multicellular systems are usually in or near metastable states corresponding to local minima, or under non-equilibrium states; therefore, our focus was not limited to thermodynamically equilibrium states corresponding to global minima. To find stable states, including local and global minima, we started our simulations from various initial particle configurations. In other words, simulation outcomes depend on not only distance–force curves but also initial configurations.

In Figure 6, two initial configurations are shown. In the case of "aggregate & cuboid" as an initial configuration, the particles have a columnar configuration with a 2×2 or 3×3 particles per layer. In the case of "aggregate & cube", the particles have a cubic configuration with 2×2×2, 3×3×3, or 6×6×6 particles per unit. In the above two cases, we added slight positional fluctuations to the initial configurations. In all four cases of the *C. elegans* and mouse embryos, the numbers of particles were set to be different so that the numbers were nearly consistent with each *in vivo* situation.

In Figure 8A and S15, two initial configurations of the particles were applied (Fig. S15A and B), and stable states were computed. In the initial configuration in Figure 8A and S15A, the particles have a columnar configuration with a 3×3 particles per layer with two layers. The particle number was set to 16, and thus there are 9 or 7 particles in the first or second layer, respectively (Fig. S15A, initial). We added slight positional fluctuations to the initial configurations. In the initial configuration in Figure S15B, the particles were nearly spherically assembled. The particle number was set to 16. The simulation results were visualized where the particle diameter was set to 25μm almost corresponding to the mean distance at the potential minimum in the EDTA-treated embryos.

In Figure 8, sphericities and aspect ratios of the simulation results were measured through generating three-dimensional (3D) images (i.e. a set of images corresponding to multiple z-slices). Each particle was depicted as a sphere with the diameter = 25μm in the 3D image. Then, the sphericities were measured by applying the MorphoLibJ plugin ("Analyze Regions 3D" menu) in Fiji/imageJ software (https://imagej.net/plugins/morpholibj). The definition of the sphericity in this plugin is as follows: sphericity = $36\pi V^2/S^3$, where $V$ or $S$ is the volume or the surface area of the object. If the particles are separated each other, the summation of the volumes or the surface areas of all particles is measured. Then, we adopted a following definition instead of the above one because this definition seems to be more generally used: sphericity = $(36\pi V^2)^{1/3}/S$. The aspect ratio was also measured by applying the MorphoLibJ plugin where ellipsoidal fitting was used. Among the three lengths of the axes in the ellipsoid, the ratio of the length of the longest axis to that of the shortest axis was calculated, resulting in the aspect ratio.

#### 6-2. Simulation data generation for systematic validation of inference method

##### 6-2-1. Force fluctuation and its persistency

We systematically generated various simulation data which were used for validating our inference method. These simulations were performed under the distance–force (DF) curves derived from the LJ potential as described previously (Section 4-3-4-1). In addition, we introduced fluctuations in the Equation S1 ($V_{C|p} = F_{C|p} / \gamma + \zeta$) (i.e. $\zeta \neq 0$). This is because, in the absence of the fluctuations, a system rapidly reaches a stable state where the particles in the system show no motions. Situations with no particle motions are not reasonable in living cellular systems. To achieve continuous motions of cells, the cells should have fluctuations of their motions. Brownian motion due to thermal fluctuations confers continuous motions, but, thermal fluctuations are negligible in cellular scale phenomena. On the other hand, cellular activities of cell–cell adhesion machineries, cytoskeletons, etc. would provide fluctuations of cellular motions with relatively long persistence time reflecting time scale of cellular processes. In the absence of external objects such as substrates, the forces generated by the cellular activities are transmitted to adjacent cells, resulting in the action-reaction relationship between the cells. So, we assumed that the effect of the fluctuations of cells are introduced into the forces of cell–cell interactions.

The forces of cell–cell interactions have been already provided from the LJ potentials, $F(D)=\frac{\kappa}{\sigma}\left\{12\left(\frac{\sigma}{D}\right)^{13}-6\left(\frac{\sigma}{D}\right)^{7}\right\}$ as a DF curve (Section 4-3-4-1). We provided the fluctuations as relative values of the forces derived from the DF curve as follow: $F_{\mathrm{wfl}}(D) = F(D) + v\,|F(D)|$, where $F_{\mathrm{wfl}}$ is the force of cell–cell interactions, wfl means "with fluctuation", and $v$ is the magnitude of the fluctuations defined as a ratio to $|F(D)|$. $v$ was defined from minus to plus values. According to this definition, because the value of $F(D)$ at the distance around the diameter of the particle is zero, the value of $v\,|F(D)|$ at the distance also becomes zero. But this is not physically reasonable, because each term of attraction and repulsion in the LJ potential at the distance around the diameter is not zero. We considered that the values of fluctuation at the distance around the diameter should be larger than or, at least, equivalent to the values at the distance providing the maximum absolute value of attractive forces. Then, we assumed a following equation:

$$F_{\mathrm{wfl}}(D) = F(D) + v(pc, \tau)\,|F(d)|, \qquad d = D, \quad \text{if } D > D_{\mathrm{max\_att}}$$
$$d = D_{\mathrm{max\_att}}, \quad \text{if } D \leq D_{\mathrm{max\_att}} \qquad \text{Eq. S9,}$$

where $D_{\mathrm{max\_att}}$ is the distance providing the maximum absolute value of attractive forces. $v$ is given by Gaussian distribution, $N(0, (pc/100)^2)$, where the average and standard deviation (SD) of the Gaussian distribution is 0 and ($pc$/100), respectively. $pc$ means the percentage. $\tau$ is the persistence time period reflecting time scale of cellular processes as mentioned in the previous paragraph. The value of $v(pc, \tau)$ is evolved for every time period $\tau$ and is unchanged during $\tau$. In the related figures, $pc$ and $\tau$ are shown as "SD value of force fluctuation (%)" and "Persistency of force fluctuation (min)", respectively.

#### 6-2-2. Simulations under various setting

##### 6-2-2-1. Assembling process

In Figure 2A, assembling processes of particles were simulated. 64 particles were initially positioned with an array of 4×4×4 cubic shape with positional fluctuations for each particle. In the absence of the force fluctuation (i.e. *pc* = 0), the particles were assembled to form a compacted configuration after the simulation time period corresponding to 1,000 min. On the other hand, in the presence of the force fluctuation, the particles were assembled, but they were not so compacted compared with the situation without the force fluctuations.

##### 6-2-2-2. Steady state

In Figures 2B, steady states of systems were simulated. The simulation processes were essentially similar to those in the assembling processes describe in Section 6-2-2-1, except that the simulations were initially run for a while (Fig. 2B, $t$ = 200 min) to reduce the influence of initial configurations on dynamics of the systems.

##### 6-2-2-3. Cell proliferation

In Figures 2C, cell proliferation/division was assumed. The time period of a cell cycle was set to be constant, and each cell undergoes cell division after the time period. At the time immediately after cell division, the distance of two daughter cells was set to be 4.3 and the centroid of the two cells was set to coincide with the position of their mother cell. The orientation of cell division was randomly searched so that distances between the daughter cells and their neighboring cells satisfy the value larger than 4.4. 1000 cycles of the random searches for the orientation were performed, and if the appropriate orientation was not found, the cell division was cancelled, and the mother cell enter a new cell cycle.

##### 6-2-2-4. Systems with spherical constraint

In Figure S4 particles were embedded into a spherical constraint. When a particle moves to the outer region of the surface of the constraint, the position of the particle was modified toward the center of the spherical constraint so that the position was just in contact with the surface (Fig. S4).

##### 6-2-2-5. Systems with spherocylindrical constraint

In Figures 3A and S5, particles were embedded into a spherocylindrical constraint whose two sides were conjugated with hemispherical constraints. When a particle moves to the outer region of the surface of the constraint, the position of the particle was modified toward the center of the hemispheres or the centerline of the spherocylinder, which is dependent on whether the particle was located in the hemispherical regions or the spherocylindrical region.

#### 7. Experimental materials and methods

##### 7-1. Mouse embryos

Embryos were obtained after mating homozygous R26-H2B-EGFP knock-in male mice which constitutively express EGFP (enhanced green fluorescent protein)-fused H2B (histone2B proteins) [17] and ICR female mice (Japan SLC). The embryos were cultured in EmbryoMax KSOM +AA with D-Glucose (Millipore, USA) covered with mineral oil on a glass bottom dish (35-mm; 27-mm ϕ, Matsunami, Japan) at 37°C, and subjected to microscopic imaging. Four distinct embryos were analyzed for each embryonic stage as shown in Figure S9.

Procedures for drug treatments are described as follows. In the case of EDTA, cytochalasin D, and blebbistatin (Fig. 7 and S11), embryos were obtained at embryonic day 2.5 (E2.5), and cultured for 12-18 hours until the embryos reached ~16 cell-stage. The embryos were transferred to medium with drugs. After ~30 min culture, the compaction states of the embryos were relaxed, and live imaging was started. After the imaging, the embryos were transferred to no-drug medium (i.e. rescue), and cultured for 1-2 days. The concentrations of EDTA, cytochalasin D, and blebbistatin are 2mM, 4μg/mL, and 100μM, respectively. Under these conditions, the embryos eventually formed the blastocysts after the rescue process, meaning that the drug treatments are not lethal to the cells and embryos. Note that we used the embryos with early phase of 16 cell-stage, because the embryos with later phase were not easy to be decompacted by the drugs and application of higher drug concentrations became lethal. Six to 10 distinct embryos were analyzed for each drug condition as shown in Figure S12.

For staining of the drug-treated embryos with anti-E-cadherin antibody or with phalloidin (Fig. 7D and S13A-B), the embryos just before the rescue process were fixed by 4% paraformaldehyde for 1 hour under r.t.. The antibody is ECCD-2, a gift from M. Takeichi [18], and phalloidin is Alexa Fluor 594-conjugated ones (A12381; invitrogen). The secondary antibody was Alexa Fluor 594 goat anti-Rat IgG (H+L) (500×dilution) (A11007; invitrogen). For staining with FM4-64 (Fig. S13C), the embryos just before the rescue process were cultured under medium with both the drugs and FM4-64 for 30 min at 37°C, and then imaging was performed. The concentration of FM4-64 (FM™4-64FX, F34653; Invitrogen) was 5ng/μL. For three-dimensional image construction (Fig. S13, "3D"), the interval of the z-sections was 1μm.

Animal care and experiments were conducted in accordance with National Institutes of Natural Sciences (NINS), the Guidelines of Animal Experimentation. The animal experiments were approved by The Institutional Animal Care and Use Committee of NINS.

#### 7-2. Microscopic live imaging

Fluorescence images were acquired on a Nikon A1 laser scanning confocal microscope (Nikon, Japan) equipped with a 20× objective (PlanApo; Dry; NA=0.75, Nikon, Japan). A stage-top $CO_2$ incubation system was used (INUG2-TIZ, Tokai Hit, Japan) on the inverted microscope. H2B-EGFP was excited using a 488-nm laser.

For imaging of mouse embryos, 50-100 frames were acquired at 3-min intervals in 30–40 z-slices separated by 2.5 μm. Nuclear tracking procedures are described in Section 2.

#### 9. Supplementary figures

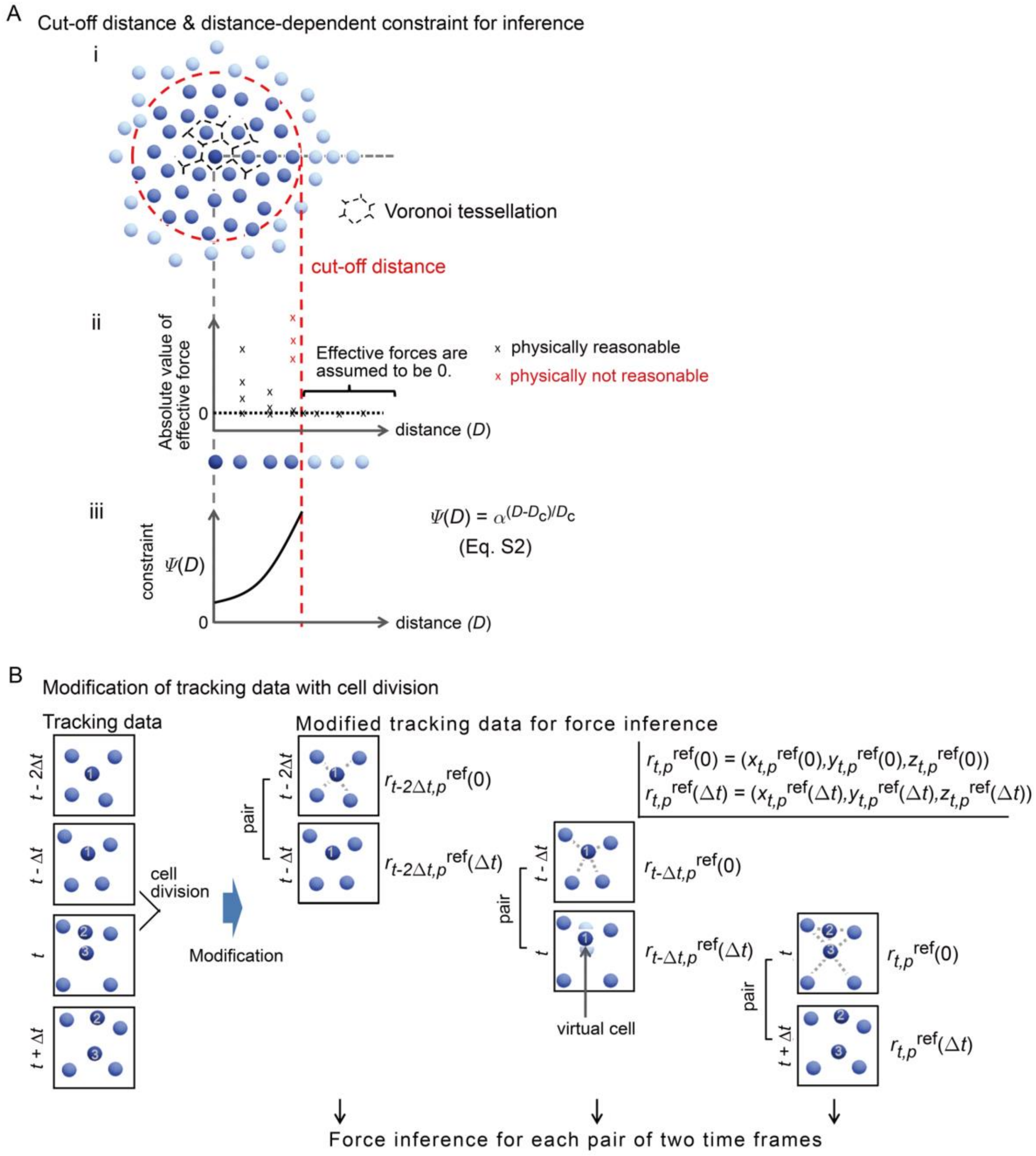


Figure S1 (related to Figure 1): Procedures to infer effective force of cell–cell interaction

A. Cut-off distance of cell–cell interaction and distance-dependent constraint for force inference. A-i. The cut-off distance is determined using the Voronoi tessellation. The Voronoi tessellation around the deepest blue particle at the center of the panel is depicted. The cut-off distance for the deepest blue particle (red broken line); particles inside the cut-off distance (deeper blue than those outside the distance). A-ii. The physically expected relationship between the distance and force. The deepest blue particle in A-i is shown. Other particles are consistent with those on the horizontal broken gray line in

A-i. In regions with a distance longer than the cut-off distance, the effective forces of cell–cell interaction are 0. To achieve physically reasonable profiles of this relationship, around regions with distances slightly shorter than the cut-off distance, the values of the effective forces should be near 0 (black crosses) and not have larger values (red crosses). A-iii. During the inference of the effective forces, to avoid physically non-reasonable values of the effective forces shown in A-ii, we introduced a distance-dependent constraint $\Psi(D)$ which is exponentially increased.

B. Nuclear tracking data with cell division is modified to be compatible with the force inference method. During time frames $(t - \Delta t)$ and $t$, a mother cell (#1) divides into two daughter cells (#2 and 3). The force inference is applied for each pair of adjacent time frames. When a cell divides, a virtual cell is assumed to exist at the centroid of the two daughter cells. Some cell–cell interactions assumed for cells #1, #2, and #3 are described by gray broken lines. The parameters in this figure are defined in Supplementary Information (Section 4-2).

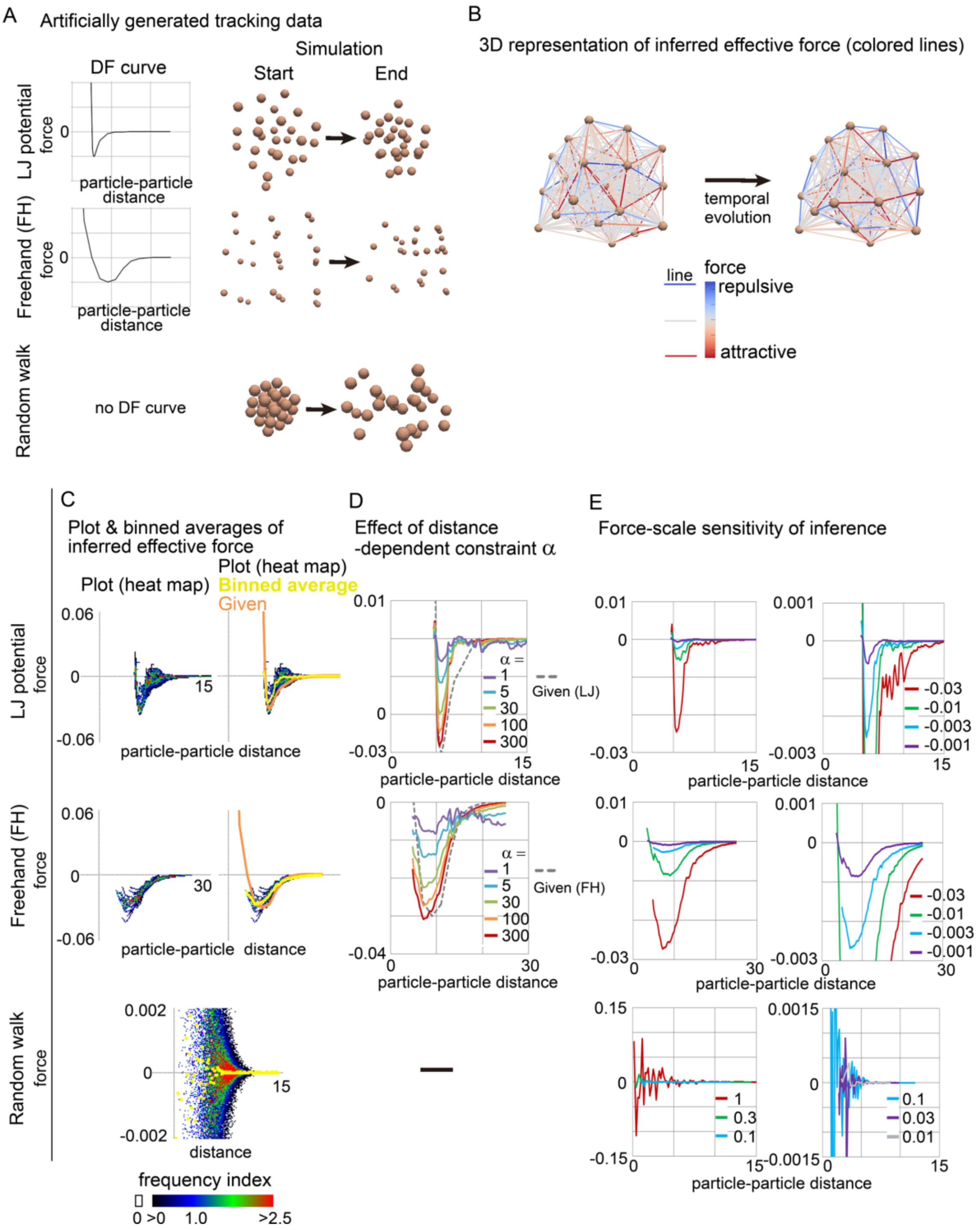


Figure S2 (related to Figure 2): Validation of force inference method using simulation data of particle tracking

A. Simulation data of particle tracking obtained from mathematical simulations of particles under the Lenard–Jones (LJ) or the freehand (FH) potential. The distance–force curves of these potentials are described. The dimensions of force and distance are arbitrary. The configurations of the particles at the starting and ending time points during the simulations are shown as light brown spheres. In addition, a random walk simulation is also shown.

B. Snapshots of outcomes of the force inference at two time frame are exemplified. Through the force inference, effective forces are obtained for each cell–cell interaction, as shown by lines colored according to the force values (red to blue). Particles, light brown.

C. Inferred effective forces from the simulations in A were plotted against the distance of cell–cell interactions. The graph space was divided into 128×128 square regions, and the frequencies of the data points plotted in each region were computed as a heat map (frequency index). Binned averages (yellow) were overlaid on the heat map (right column). The original distance–force curves defined in A are also merged (orange; given) in the case of the LJ and FH potentials.

D. Effect of distance-dependent weight defined in Equation S4 on inference results. The binned average data obtained in C are used as distance–force curves. In the absence of the distance-dependent constraint ($\alpha$ = 1), the inferred distance–force curves are close to 0 (purple lines) and are far from the given profiles (gray broken lines) for both the LJ and FH potentials. When the distance-dependent weight increases, the inferred distance–force curves approach the given profiles ($\alpha$ = 100, 300).

E. Sensitivity of the inference method to force scales. LJ and FH potentials were prepared with various minima of the force values, simulations were performed, and effective forces were inferred. Then, distance–force curves were estimated. The provided minima are shown in the figures (-0.03, -0.01, -0.003, and -0.001). In the case of the random walk, simulations under various magnitudes of random movements were provided as shown as relative values (1, 0.3, 0.1, 0.03, and 0.01), and the distance–force curves were randomized.

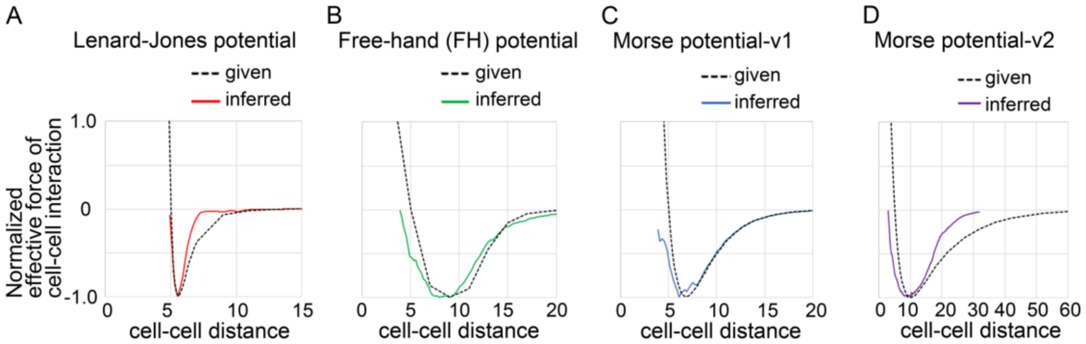


Figure S3 (related to Figure 2): Validation of force inference method for various potentials

Distance–force (DF) curves were inferred from simulation data generated under various potentials. The force values were normalized by the maximum attractive forces in the given potential. The mean cell diameters were set to be 5.0, where the forces are 0. The given potentials are the LJ (A), FH (B), and Morse (C and D) from previous two papers [19,20]. Solid lines, inferred DF curves; broken lines, the given potentials.

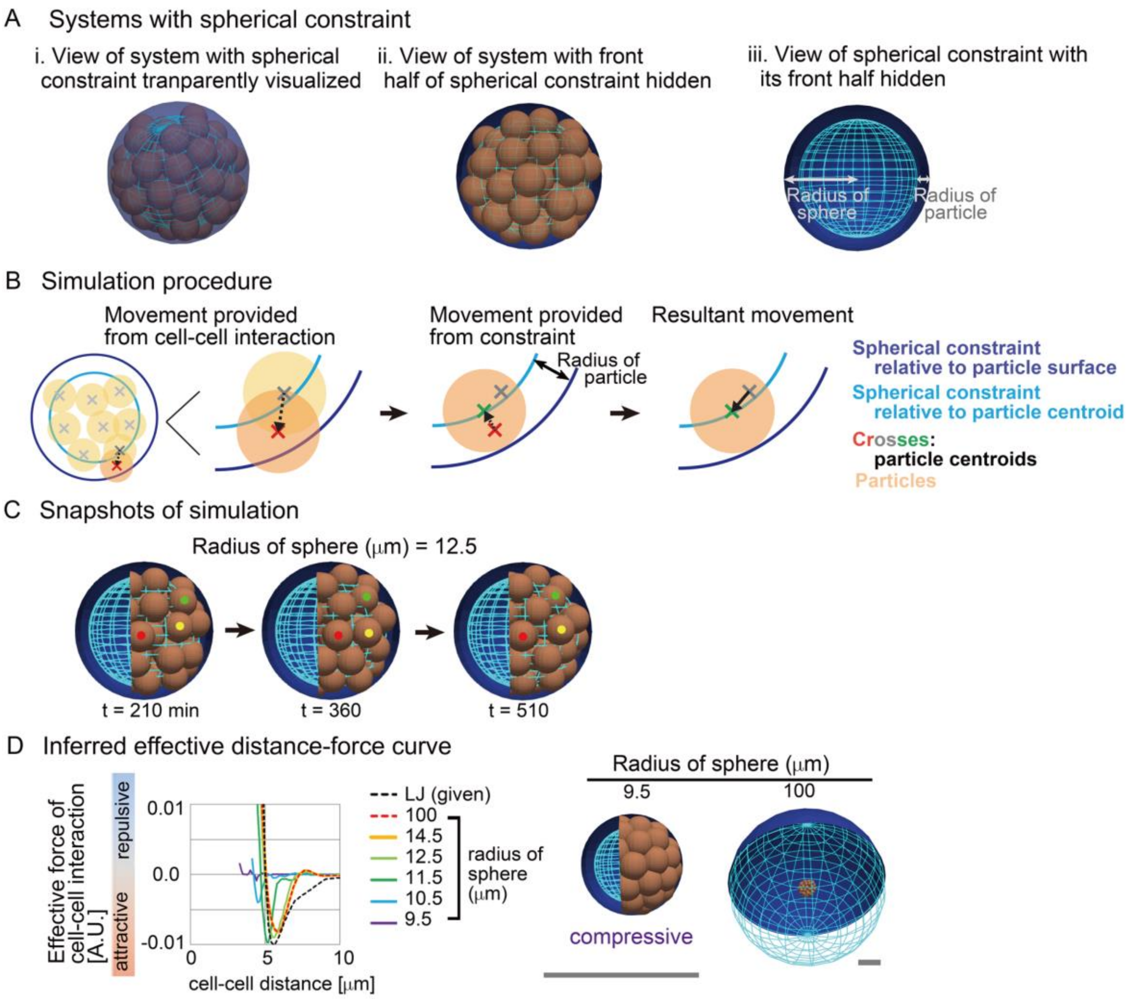


Figure S4 (related to Figure 3): Inference of effective forces under spherical constraints

Simulation data were used to validate our inference method. Systems with spherical constraints were considered.

A. Particles were embedded into a spherical constraint (A-i and -ii). Definitions of the radius of the sphere and the radius of particles are shown (A-iii).

B. The simulation procedures when a particle collides with the spherical constraint are described (dark orange circle). The detailed procedures are described in Supplementary Information (Section 6-2-2).

C. Snapshots of simulations. The Lenard-Jones potential (LJ) was provided. Three particles are marked by red, green, and yellow. The number of particles were set to be 64 in all conditions. The condition is [sampling interval = 3.0 min, SD value of force fluctuation = 1000, and persistency of force fluctuation = 1 min].

D. Inferred DF curves under different radius of spheres. Two snapshots under the different radius are shown. In the case that 9.5μm, the particles were very closely contacted each other so that the distances between the adjacent particles seemed to be less than the diameter of the particles. Therefore, each particle was compressed. In the case that the radius was 100μm, the spherical constraint was sufficiently large so that the particles are not in contact with the surface of the constraint. The black scale bars and the gray ones correspond to the diameter of the particles and the diameter of the spherical constraint with the radius = 14.5μm.

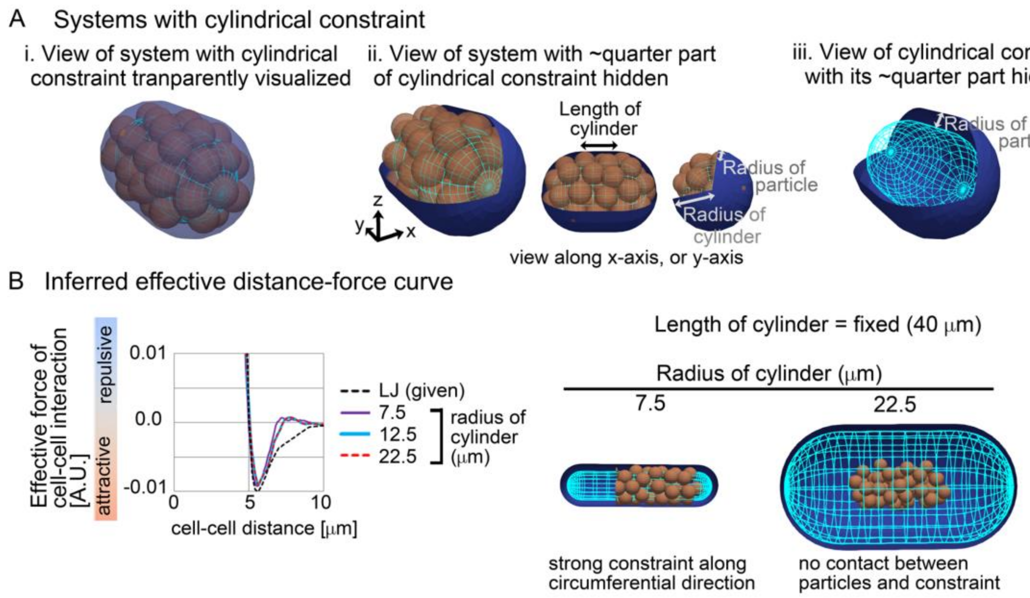


Figure S5 (related to Figure 3): Inference of effective forces under spherocylindrical constraints

Simulation data were used to validate our inference method. Systems with spherocylindrical constraints were implemented.

A. Particles were embedded into a spherocylindrical constraint (A-i and -ii). Definitions of the radius and the length of the spherocylinder, and the radius of particles are shown (A-ii and -iii). The simulation procedures when a particle collides with the spherocylindrical constraint were implemented in a similar manner to Figure S4.

B. Inferred DF curves under different radius of the spherocylindrical constraints. Two snapshots under the different radius are shown. The particles are in close contact with the surface of the constraint in the case that the radius was 7.5μm. In the case that the radius was 22.5μm, the spherocylindrical constraint was sufficiently large so that the particles are not in contact with the surface of the constraint. The number of particles were set to be 54 in all conditions. The condition is [sampling interval = 3.0 min, SD value of force fluctuation = 1000, and persistency of force fluctuation = 1 min].

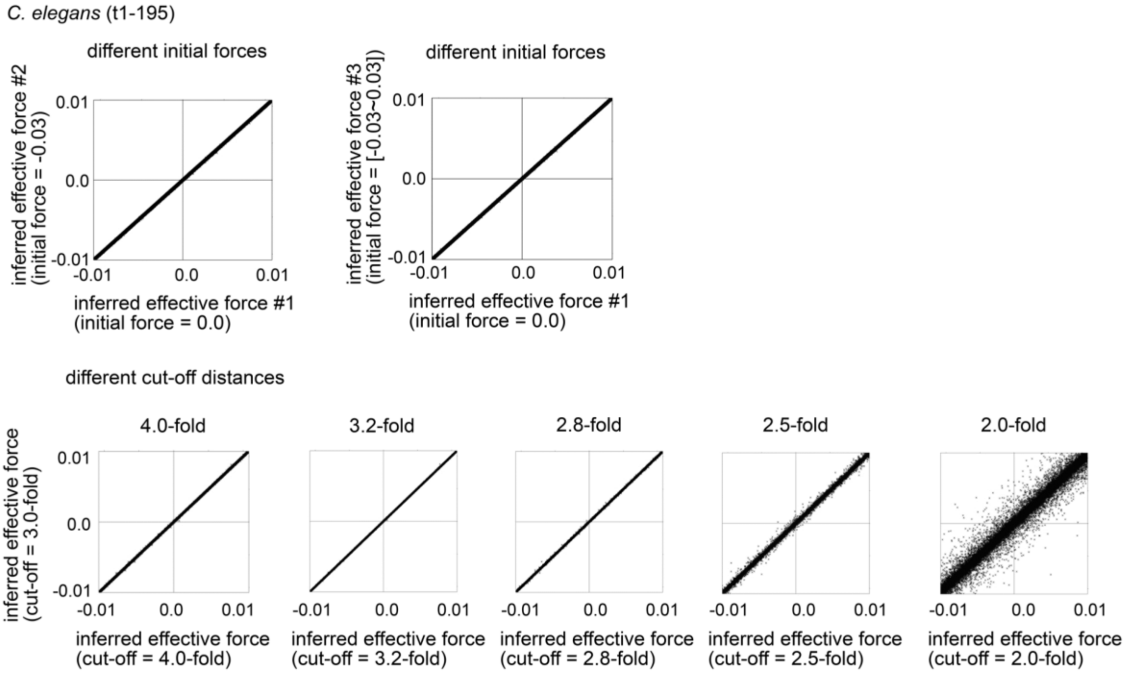


Figure S6 (related to Figure 4): Uniqueness of solutions of effective force inference in *C. elegans*

In the top left panel, the minimizations of Equation S6 were performed from different initial force values as described in the x- and y- axes. In the top right panel, the initial forces were given as uniform random numbers ranging from -0.03 to 0.03 in the y-axis. The inferred values of each cell-cell interaction were plotted by crosses. The inferred values from the different initial force values were absolutely correlated in the all cases, suggesting that a unique solution was obtained in each system. In the bottom panels, different cut-off distances were given as indicated (3.0, 4.0, 3.2, 2.8, 2.5 and 2.0-fold diameter of cell bodies), and the minimizations were performed. The inferred values were strongly correlated when the cut-off distances were greater than 2.8-fold diameter of cell bodies, suggesting that a unique solution was obtained under these conditions.

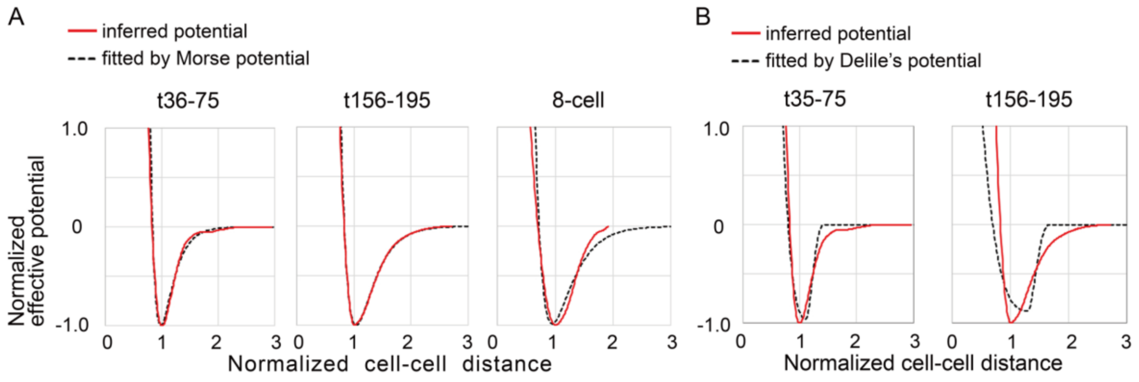


Figure S7 (related to Figure 4): Fitting of the previously-reported potentials to inferred distance–potential (DP) curves in *C. elegans* and mouse embryos

A. The Morse potential was used for fitting. The formula of the Morse potential is: $U(D) = U_e\,[\exp\{-2a(D-D_e)\} - 2\exp\{-a(D-D_e)\}]$, where $U$ is the potential energy, $D$ is the particle–particle distance, and $U_e$, $D_e$, and $a$ are the fitting parameters. Cell–cell distances were normalized by the distances providing the potential minima in the inferred potentials. Effective potential energies were normalized so that the potential minima become -1.0. t36-75 and t156-195 are from the *C. elegans* as defined in Fig. 4, and 8-cell is from the mouse 8-cell stage embryo. Fitting was performed by using the solver implemented in the Excel software.

B. A potential proposed by Delile et al. was used for fitting [20]. The distance–force curve was described in the Delile's paper, from which we numerically computed the DP curve for fitting.

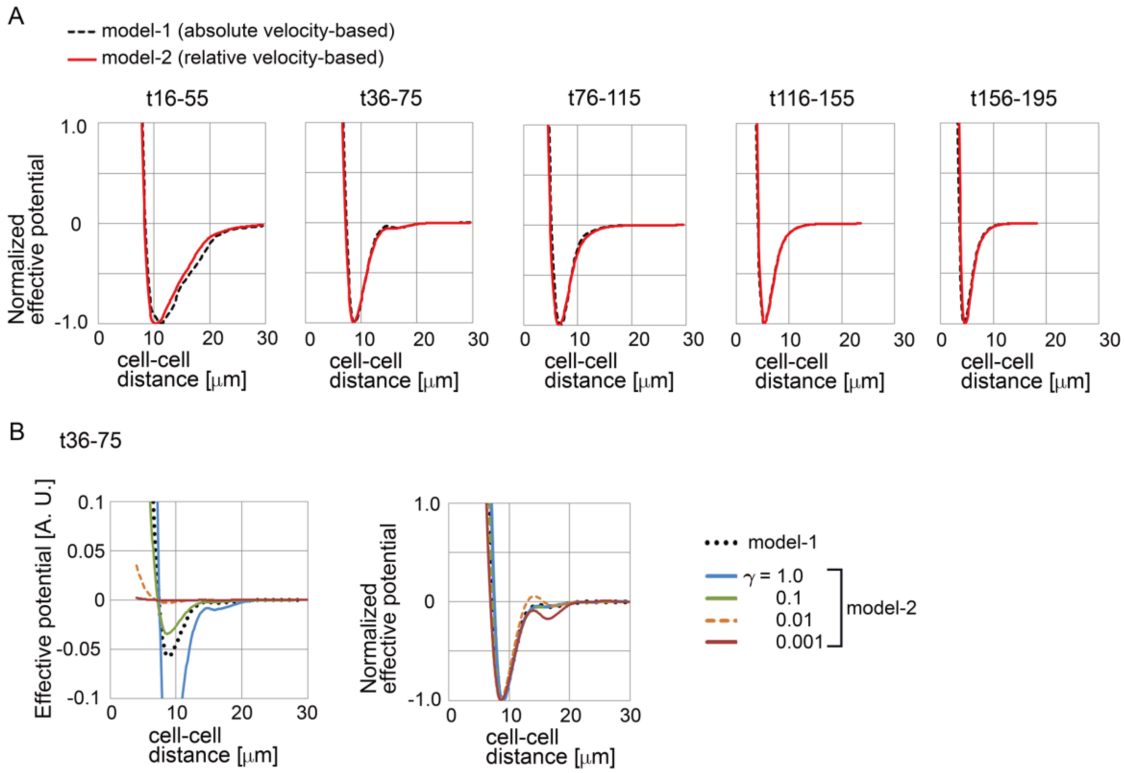


Figure S8 (related to Figure 4): Inferred distance–potential (DP) curves under the assumption of the relative velocity-dependent model (Eq. S1-2) in *C. elegans* embryos

A. Comparison of the inferred DP curves under the two models: the absolute velocity-based and the relative velocity-based models. The effective potential energies were normalized so that the potential minima become -1.0. The time frames (e.g., t16-55, etc) were defined in Fig. 4.

B. Parameter dependency of inferred DP curves in *C. elegans* embryo at the t36-75 time frame. The values of $\gamma$ in Equation S1-2 were variously set. The inferred DP curve from the absolute velocity-based model (model-1) is also presented for comparison, where $\gamma$ was set 1.0. In the relative velocity-based model, because $\omega_g(D_{pm})$ in the Equation S2-1 was set to be 1.0 at $D_{pm}$ = the diameters of cell bodies, the values of $\{\omega_g(D_{pm})\,\gamma\}$ at $D_{pm}$ = the diameters of cell bodies are equal to $\gamma$.

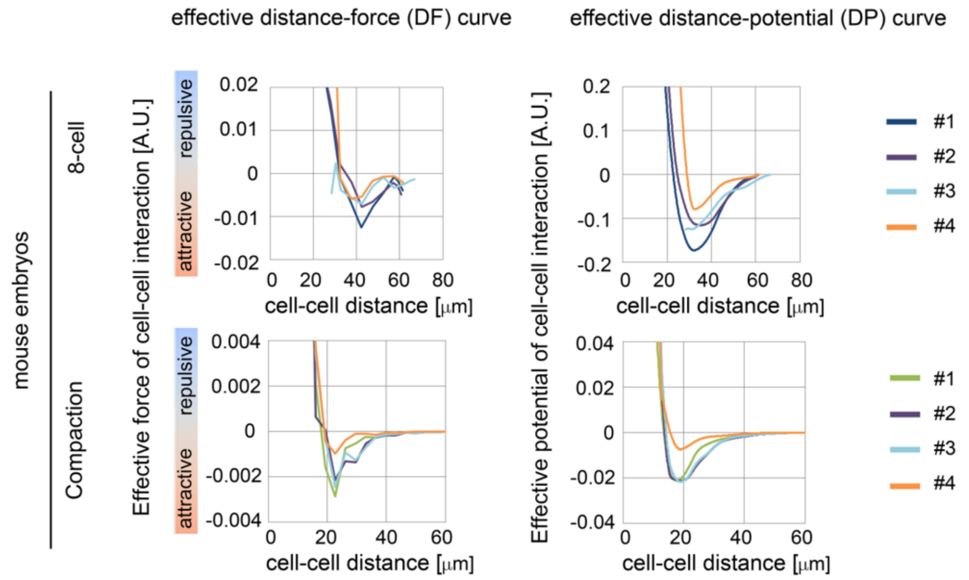


Figure S9 (related to Figure 5): Distance–force and distance–potential curves in mouse embryos

Inferred DF and DP curves in mouse 8-cell and compaction stages. Four independent embryos (#1-4) were analyzed for each stage. #1 for each stage corresponds to Figure 5. Uniqueness of solutions was confirmed by a similar way to Fig. S6 (data not shown).

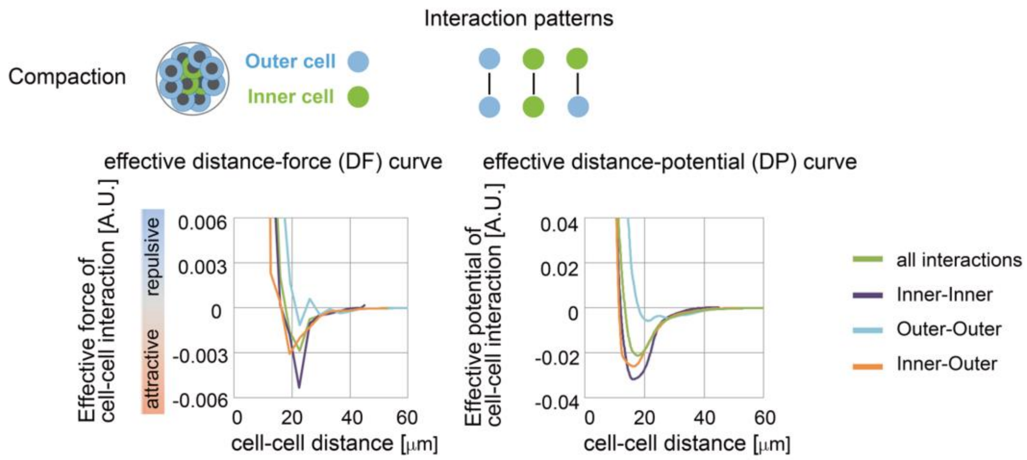


Figure S10 (related Figure 5): Distance–force and distance–potential curves of different cell types in mouse embryos

A. Inferred DF and DP curves of outer (blue circles) and inner (green circles) cells in mouse compaction stage. There are three possible interactions: inner–inner, outer–outer, and inner–outer cells. These data were obtained from embryo #1 in Figure S9, compaction. The curves of all interactions (green lines) are identical to those in Figure S9.

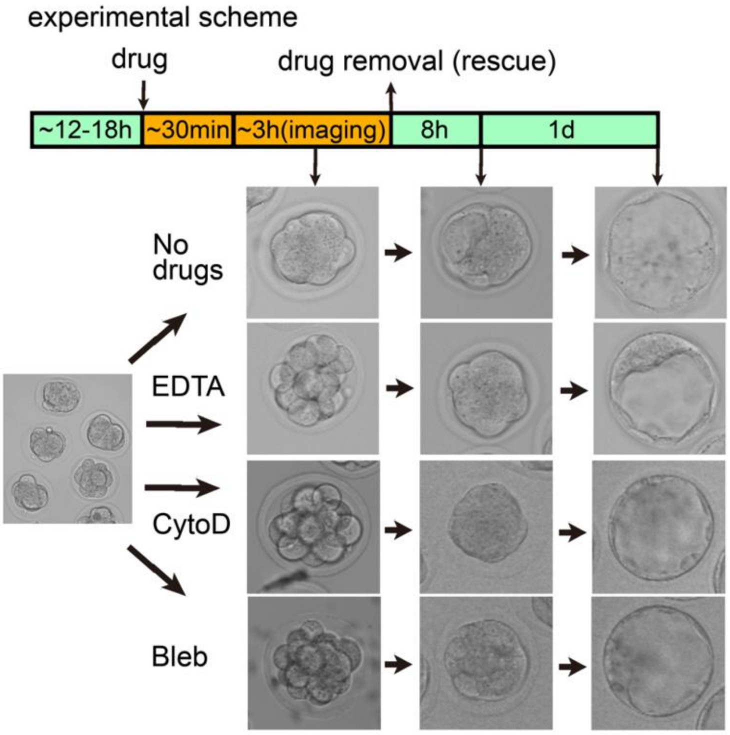


Figure S11 (related to Figure 7): Experimental design for inhibiting compaction in mouse embryos

The experimental design of Figure 7 is shown. The upper panel is the experimental scheme. The lower panels are microscopic images of embryos at each step of the experimental scheme. CytoD, cytochalasin D; Bleb, blebbistatin.

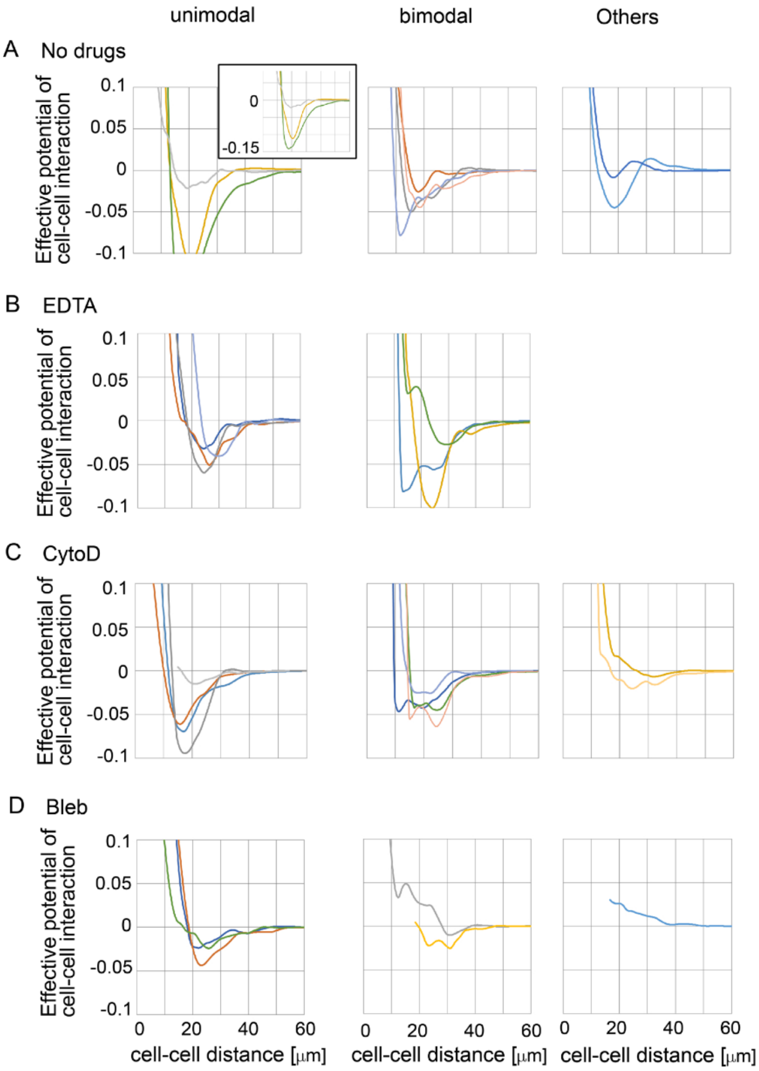


Figure S12 (related to Figure 7): Distance–potential curve in compaction-inhibited mouse embryos

The DP curves in Figure 7 are enlarged. For visualization, the DP curves were roughly categorized into three groups; unimodal (almost single potential minimum), bimodal (almost double potential minima), and others (the distance at potential minimum is very long, or a potential maximum exists). CytoD, cytochalasin D; Bleb, blebbistatin.

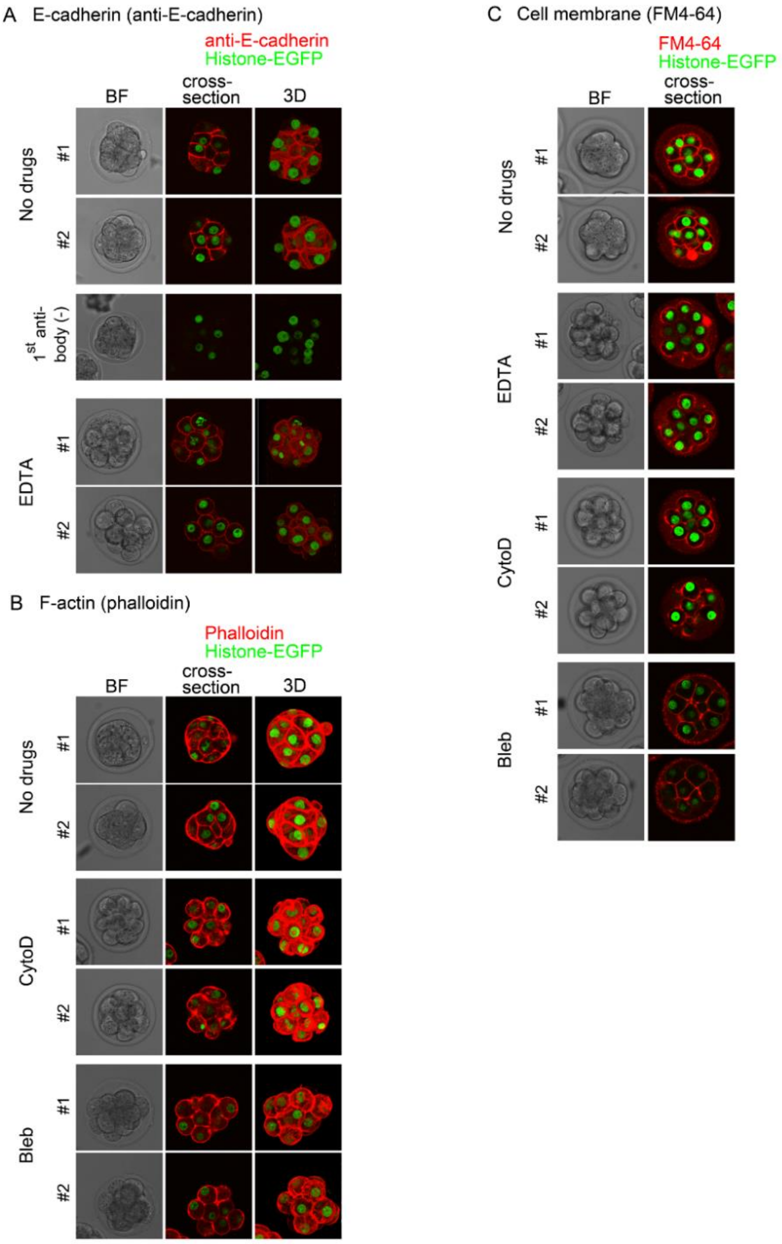


Figure S13 (related to Figure 7): Histological observations of compaction-inhibited mouse embryos

A. Immuno-staining of E-cadherin. Two examples are shown for the normal or EDTA-treated embryos (#1 and #2). Conditions without the 1st anti-body is also shown as the negative control. BF, bright field; 3D, three-dimensional image. Green, Histone-EGFP.

B. Phalloidin staining for F-actin. Two examples are shown for the normal, cytochalasin D-treated, or blebbistatin-treated embryos (#1 and #2). BF, bright field; 3D, three-dimensional image. Green, Histone-EGFP. CytoD. Cytochalasin D; Bleb, blebbistatin.

C. FM4-64 staining for cell membrane. Two examples are shown for the normal, EDTA-treated, cytochalasin D-treated, or blebbistatin-treated embryos (#1 and #2).

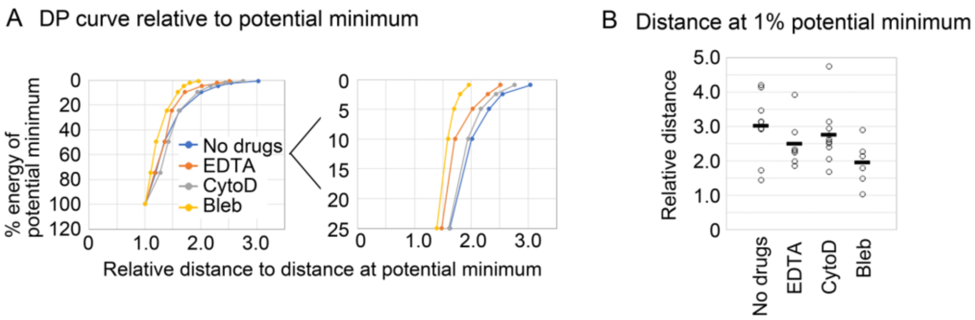


Figure S14 (related to Figure 7): Quantitative comparison of profiles of distance–potential curves in compaction-inhibited mouse embryos

A. Related to the right panel of Figure 7C. Distances at given % of potential minima were calculated as the relative distances to the distances at the potential minima. 75, 50, 25, 10, 5, 2.5, and 1 % were considered. The mean values were plotted. CytoD, cytochalasin D; Bleb, blebbistatin.

B. The results under 1% in A. Mann–Whitney–Wilcoxon tests were performed and the resultant *p*-values for "No Drugs" vs. "EDTA", vs. "CytoD", and vs. "Bleb" are 0.21, 0.21, and 0.036, respectively. In the case for 10% potential minima (Fig. 7C, right panel), the *p*-values are 0.30, 0.60, and 0.11. Black bar, mean.

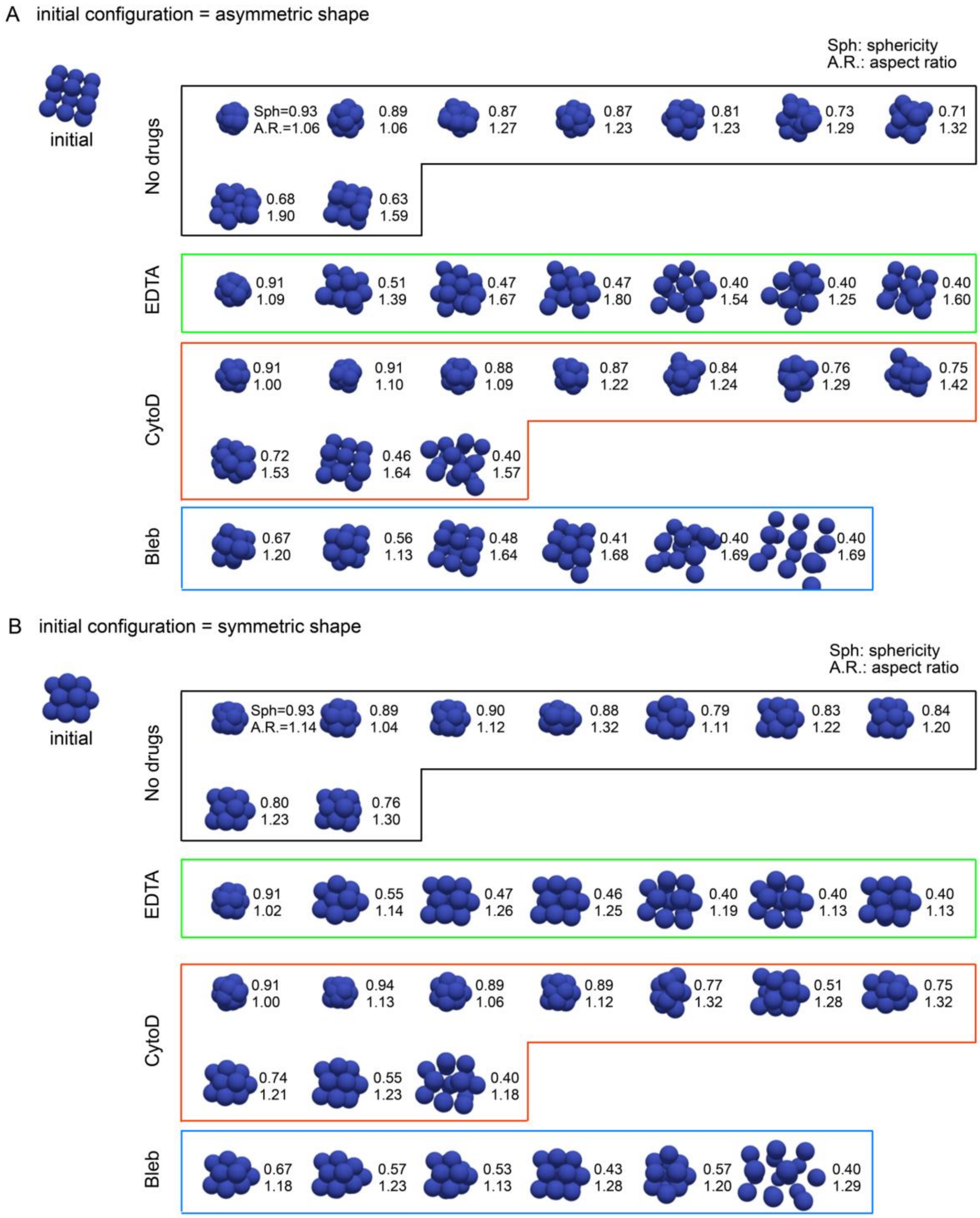


Figure S15 (related to Figure 8): List of simulation results of compaction-inhibited mouse embryos

Simulations were performed under the all DP curves obtained from the compaction-inhibited embryos in Figure S12.

A. The initial configuration of particles was given as an asymmetric (not symmetric nor spherical) shape (initial). The simulation results were ordered according to the sphericities. The values of the sphericities and of the aspect ratios were shown. Sph, sphericity; A.R., aspect ratio. The percentages of final shapes with sphericity > 0.8 are 56 (no drugs), 14 (EDTA), 50 (cytochalasin D), or 0% (blebbistatin), respectively. The sphericities and aspect ratios are plotted in Figure 8B and C.

B. The initial configuration of particles was given as a nearly symmetric/spherical shape (initial), and the similar analyses to A were performed. The order of the simulation results was set to be the same as that in A.

#### 10. Video

Uploaded on https://doi.org/10.6084/m9.figshare.21714842

Figure 4-video 1 (Movie S1A): 3D cell movement; *C. elegans* embryo

This movie corresponds to Figure 4A, panel "Cell tracking".

Figure 4-video 2 (Movie S1B): 3D force map of cell–cell interaction; *C. elegans* embryo

This movie corresponds to Figure 4C, panel "3D representation of effective force".

Figure 5-video 1 (Movie S2A): Live imaging of three-dimensional systems; mouse 8-cell stage

Figure 5-video 2 (Movie S2B): Live imaging of three-dimensional systems; mouse compaction stage

Movies S2A-B correspond to Figure 8A, panel "Tracking".

Figure 5-video 3 (Movie S2C): 3D force map of cell–cell interaction; mouse 8-cell stage

Figure 5-video 4 (Movie S2D): 3D force map of cell–cell interaction; mouse compaction stage

Movies S2C-D correspond to Figure 5B.

#### 11. Source data

Nuclear tracking data

Uploaded on https://doi.org/10.6084/m9.figshare.21714842

Figure 5-Source Data 1 contains following files.

Nuclear tracking data; mouse 8-cell stage (sample1-4.csv)

Nuclear tracking data; mouse compaction stage (sample1-4.csv)

3D force data

Uploaded on https://doi.org/10.6084/m9.figshare.21714842

Figure 4-Source Data 1 contains following files.

3D force data of cell–cell interaction; *C. elegans* embryo (1 sample: out_force_data.dat, out_xyz_data.dat)

Figure 5-Source Data 2 contains following files.

3D force data of cell–cell interaction; mouse 8-cell stage (4 samples: out_force_data.dat, out_xyz_data.dat)

3D force data of cell–cell interaction; mouse compaction stage (4 samples: out_force_data.dat, out_xyz_data.dat)

3D force map can be visualized by reading following files in the ParaView software (https://www.paraview.org/). These files will be uploaded on Web sites.

out_display_for_ParaView_line_ref[N].vtk

out_display_for_ParaView_xyz_sphere_polygon_ref[N].vtk

Data of distance–force/potential profiles

Uploaded on https://doi.org/10.6084/m9.figshare.21714842

Figure 6-Source Data 1 contains following files.

Distance–force/potential profiles; C. elegans embryos (DF_DP_Celegans.xlsx)

Distance–force/potential profiles; mouse 8-cell stage (DF_DP_mouse_8_cell_stage.xlsx)

Distance–force/potential profiles; mouse compaction stage (DF_DP_mouse_compaction_stage.xlsx)

Source codes

Uploaded on GitHub: https://doi.org/10.5281/zenodo.7427050
